# *Haloferax volcanii* immersed liquid biofilms develop independently of known biofilm machineries and exhibit rapid honeycomb pattern formation

**DOI:** 10.1101/2020.07.18.206797

**Authors:** Heather Schiller, Stefan Schulze, Zuha Mutan, Charlotte de Vaulx, Catalina Runcie, Jessica Schwartz, Theopi Rados, Alexandre W. Bisson Filho, Mechthild Pohlschroder

## Abstract

The ability to form biofilms is shared by many microorganisms, including archaea. Cells in a biofilm are encased in extracellular polymeric substances that typically include polysaccharides, proteins, and extracellular DNA, conferring protection while providing a structure that allows for optimal nutrient flow. In many bacteria, flagella and evolutionarily conserved type IV pili are required for the formation of biofilms on solid surfaces or floating at the air-liquid interface of liquid media. Similarly, in many archaea it has been demonstrated that type IV pili and, in a subset of these species, archaella are required for biofilm formation on solid surfaces. Additionally, in the model archaeon *Haloferax volcanii*, chemotaxis and AglB-dependent glycosylation play important roles in this process. *H. volcanii* also forms immersed biofilms in liquid cultures poured into Petri dishes. This study reveals that mutants of this haloarchaeon that interfere with the biosynthesis of type IV pili or archaella, as well as a chemotaxis-targeting transposon and *aglB*-deletion mutants, lack obvious defects in biofilms formed in liquid cultures. Strikingly, we have observed that these liquid-based biofilms are capable of rearrangement into honeycomb-like patterns that rapidly form upon removal of the Petri dish lid, a phenomenon that is not dependent on changes in light or oxygen concentration but can be induced by controlled reduction of humidity. Taken together, this study demonstrates that *H. volcanii* requires novel, as-yet-unidentified strategies for immersed liquid biofilm formation and also exhibits rapid structural rearrangements.

**Importance:** This first molecular biological study of archaeal immersed liquid biofilms advances our basic-biological understanding of the model archaeon *Haloferax volcanii*. Data gleaned from this study also provide an invaluable foundation for future studies to uncover components required for immersed liquid biofilms in this haloarchaeon and also potentially for liquid biofilm formation in general, which is poorly understood compared to the formation of biofilms on surfaces. Moreover, this first description of rapid honeycomb pattern formation is likely to yield novel insights into the underlying structural architecture of extracellular polymeric substances and cells within immersed liquid biofilms.

## Introduction

Prokaryotes have evolved a variety of strategies to mitigate the effects of environmental stress, including the establishment of biofilms, which are complex microbial communities surrounded by a matrix of extracellular polymeric substances (EPS). Of the bacteria and archaea found above the subsurface, an estimated 80% in soil and upper oceanic sediment exist in biofilms (1). It has been suggested that life within a biofilm may be the primary way of active life for both bacterial and archaeal species (1), with other bacterial-specific studies suggesting that life in a biofilm may be the default, with planktonic cells merely serving as mediators for the transition from one biofilm to the next (2). The advantages of being within a biofilm for bacterial cells range from communication and environmental stress protection to improved nutrient acquisition (3). Similarly, for archaeal species, the demonstrated advantages of living in a biofilm include conferring environmental stress protection, horizontal gene transfer, and syntrophy facilitation as well as mechanical and structural stability provided by EPS (4–7). While some biofilms, such as those that play roles in wastewater treatment or bioremediation (8, 9) can provide a variety of important benefits to humans, others can cause serious harm, such as debilitating chronic infections (10–12), as biofilms confer reduced antibiotic and antimicrobial sensitivity (13, 14) that can render the embedded bacterial cells up to 1000 times less susceptible to treatments relative to planktonic cells (15). Thus, understanding biofilm formation is of significant public health interest.

A variety of proteins necessary for biofilm formation have been identified and characterized in an array of bacterial species. Biofilm formation requires type IV pili in organisms such as *Pseudomonas aeruginosa* and *Vibrio cholerae* (16–21). Flagella are also sometimes required for biofilms, such as those of *Escherichia coli* and *P. aeruginosa* under certain conditions (18, 19, 22, 23). Additionally, in *P. fluorescens*, various surface adhesins are often critical to this process (24–26). While biofilms forming at surfaces have been extensively studied, much less is known about biofilms that form in liquid media. *Bacillus subtilis* and *P. aeruginosa*, for example, form pellicles — a type of biofilm that floats at the air-liquid interface of a culture — and flagella-based motility is important for successful pellicle formation in both organisms (27–29). Chemotaxis and oxygen sensing have also been shown to play crucial roles in the formation of pellicles in *B. subtilis* (29) and quorum sensing, a form of cell-cell communication, has been shown to be required for proper biofilm formation in species such as *V. cholerae* through the regulation of EPS biosynthesis (30). Cellular appendages as well as EPS are also determining factors in shaping the structure of biofilms through cell-cell and cell-surface interactions. The involved physico-mechanical forces can range from adsorption/adhesion (coil formation, bridging), often via type IV pili, to repulsion-driven depletion attraction (phase separation), e.g. via EPS (31–34). Beyond cellular components, environmental conditions can also play a role in influencing biofilms, such as relative humidity (RH) levels and temperature (35).

Archaea also readily form biofilms in a variety of habitats (7). The genetically tractable cren- and euryarchaeal species tested thus far form surface-attached biofilms in a type IV pilus-dependent manner, and, in a subset of these species (such as *Sulfolobus acidocaldarius* and *Methanococcus maripaludis*), biofilm formation is also dependent on the archaella — structures analogous to the bacterial flagella — under certain conditions (7, 36, 37). The model haloarchaeon *Haloferax volcanii* can form biofilms on surfaces at the air-liquid interface (ALI) of a culture in a type IV pili-dependent but archaella-independent manner (38). Strains lacking the genes encoding the adhesion pilins, the prepilin peptidase, or components of the pilus biosynthesis pathway (Δ*pilA1-6, ΔpibD*, and *ΔpilB1C1* or *ΔpilB3C3*, respectively) are impaired in adhesion to coverslips at the ALI (36, 38–41). While biofilm formation in *H. volcanii* presumably also requires the chemotaxis machinery, as transposon insertions between the *cheB* and *cheW1* genes result in a mutant having an adhesion defect, *H. volcanii* biofilm formation is not impaired in a non-motile mutant lacking the archaellins *arlA1* and *arlA2* (38, 42).

Archaea can also be found in floating liquid biofilms (43, 44). Moreover, Chimileski et al. recently described *H. volcanii* immersed liquid biofilms that form in static-liquid cultures (6). These biofilms contain polysaccharide, based on Concanavalin A staining, and eDNA, based on DAPI staining, as major structural components, and possibly also include amyloid proteins based on Congo Red and Thioflavin T staining (6). Chimileski et al. also reported that after homogenization of the immersed liquid biofilm, aggregation occurred in as little as three hours, and the biofilm became more concentrated and denser over the course of 48 hours (6). However, the molecular mechanisms required for the formation of these biofilms are not yet known.

Here we report that *H. volcanii* immersed liquid biofilms form independently of type IV pili along with chemotaxis and archaella machineries, demonstrating that the mechanisms required for the formation of *H. volcanii* immersed liquid biofilms differ significantly from those required for the formation of an archaeal biofilm on an abiotic surface. We also for the first time describe a unique, rapid change in the macroscopic, three-dimensional organization of the biofilm, forming a honeycomb-like pattern in response to reduction in humidity levels, potentially revealing a strategy to disperse from a biofilm.

## Materials and Methods

### Strains and growth conditions

*H. volcanii* wild-type strain H53 and its derivatives (Table 1) were grown aerobically at 45°C in liquid (orbital shaker at 250 rpm) or on solid semi-defined Hv-Cab medium (45). H53, Δ*pibD*, Δ*pilA1-6*, Δ*pilB1C1*, Δ*pilB3C3*, Δ*pilB1C1B3C3*, Δ*arlA1*, Δ*arlA2*, Δ*arlA1-2*, Δ*aglB*, and Δ*agl15* media were additionally supplemented with tryptophan and uracil (both at 50 µg · ml^-1^ final concentration); *cheB::tn, cheF::tn*, Δ*pssA*, and Δ*pssD* media were supplemented with uracil (50 µg · ml^-1^ final concentration); H98 and Δ*cetZ1* media were supplemented with thymidine and hypoxanthine (both at 40 µg · ml^-1^ final concentration) as well as uracil (50 µg · ml^-1^ final concentration) (46). Solid media plates contained 1.5% (wt/vol) agar. *Haloferax mediterranei* was grown aerobically at 45°C in Hv-Cab medium (45).

**Table 1:**
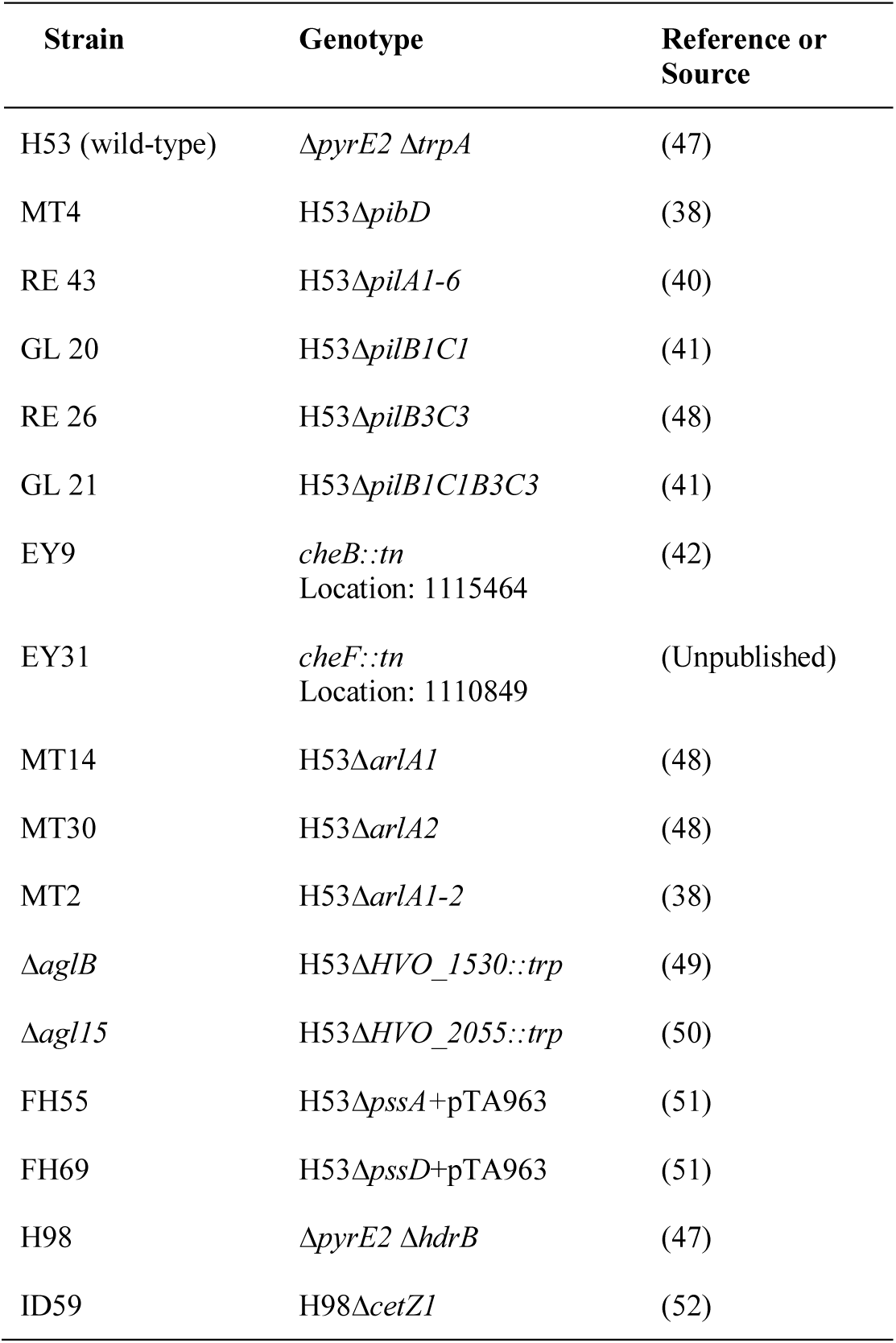
Strains used in this study.

### Immersed liquid biofilm formation

Biofilms of strains tested in this study were prepared and observed as follows: strains were inoculated in 5 mL of Hv-Cab medium followed by overnight incubation at 45°C with shaking (orbital shaker at 250 rpm) until the strains reached mid-log phase (OD_600_ 0.3-0.7). Mid-log cultures were diluted to an OD_600_ of 0.05 at a final volume of 20 mL followed by shaking incubation at 45°C for 48 hours. Cultures were poured into sterile Petri dishes (100mm x 15mm, Fisherbrand) after the 48-hour incubation period. Poured cultures were placed in plastic containers and incubated at 45°C without shaking for 18 ± 3 hours after which the resulting immersed liquid biofilms were observed and imaged.

### Honeycomb pattern observation

After an incubation period of 18 ± 3 hours post-pouring with resulting immersed liquid biofilm formation, *H. volcanii* wild-type and mutant strain cultures were observed for honeycomb pattern formation. Without disturbing the biofilms, the lids of the Petri dishes were removed immediately after plastic container lid removal and observations were made on the speed and formation of the honeycomb pattern along with its dispersal. Honeycomb pattern formation was recorded and/or imaged using an iPhone (Fig. 1, Fig. 4, Fig. S2, and Fig. S4), Canon EOS Digital Rebel XSi (Fig. 3), and Nikon D3500 DX-Format DSLR Two Lens (lens 18-55mm f/3.5-5.6G, video setting 60fps) (Fig. 2, Fig. S5, Movie S1, and Movie S2).

**Figure 1:**
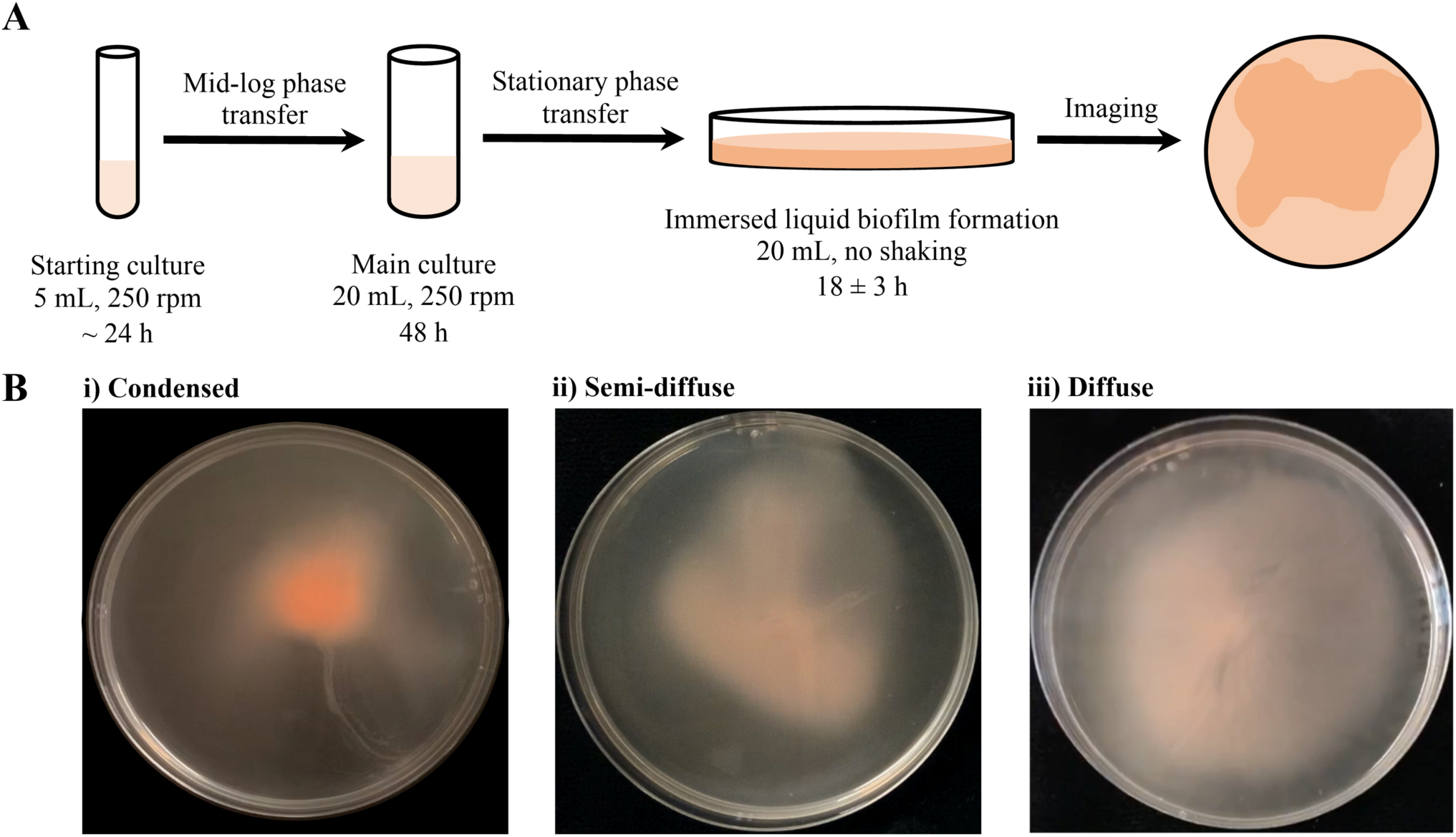
Optimized protocol for *H. volcanii* immersed liquid biofilm formation. **(A)** A schematic description of the protocol used for the reproducible observation of immersed liquid biofilm formation is shown. Single colonies are inoculated and incubated while shaking until they reach mid-log phase (OD_600_ between 0.3 and 0.7). Cultures are then diluted to an OD_600_ of 0.05, to ensure the same starting OD_600_ for different cultures, and incubated again on a shaker for 48 hours, at which point they are in stationary phase (OD_600_ of 1.8 or greater). The cultures are then poured into sterile plastic Petri dishes and statically incubated. Immersed liquid biofilm formation can be observed reproducibly after 18 ± 3 hours. All incubations were performed at 45°C. **(B)** Representative images for stochastic variations in the shape and color of immersed liquid biofilms for the wild-type are shown, ranging from dark, condensed (i) to light, diffuse immersed liquid biofilms (iii). All immersed liquid biofilms are imaged after 18 ± 3 hours of static incubation at 45°C. The diameter of the Petri dishes is 10 cm.

**Figure 2:**
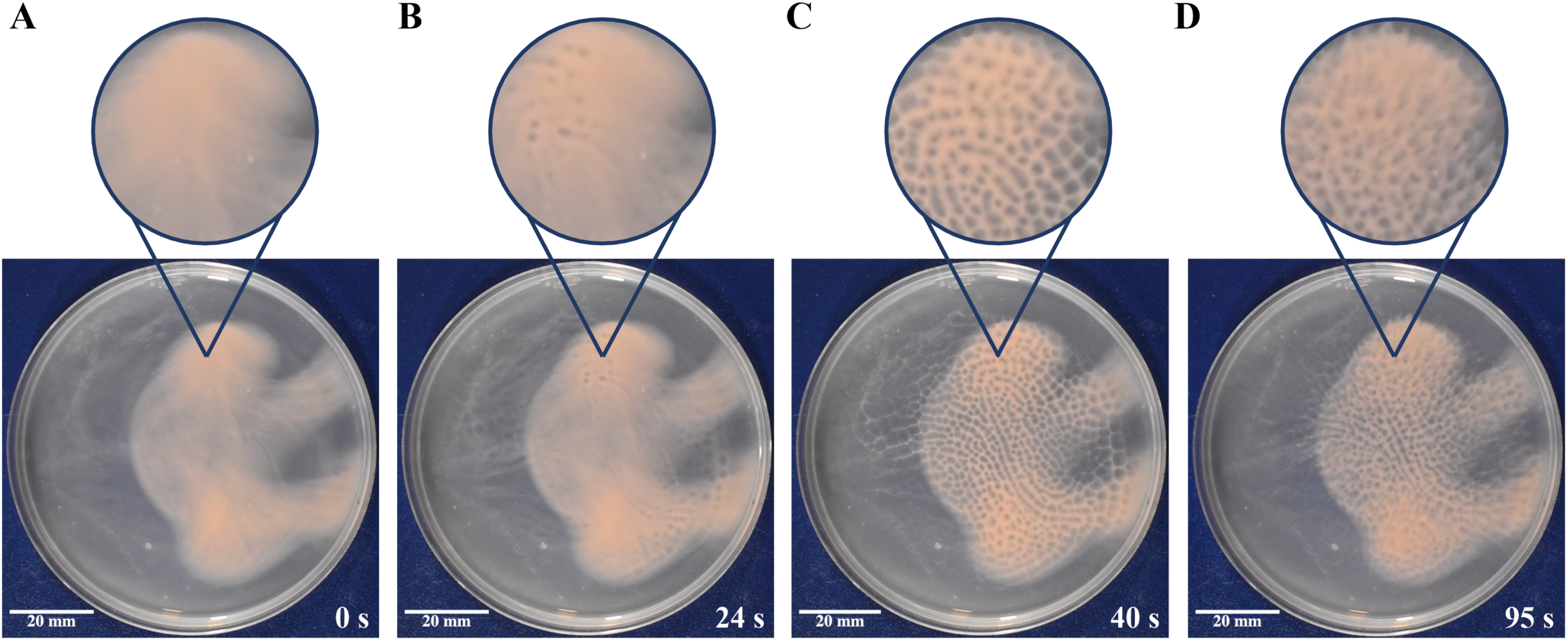
Immersed liquid biofilms exhibit honeycomb pattern formation. Prior to the experiment, wild-type immersed liquid biofilms had formed after 18 h incubation at 45°C. **(A)** Images were taken immediately after Petri dish lid removal, followed by **(B)** start of honeycomb formation 24 seconds after lid removal, **(C)** peak honeycomb pattern formation 40 seconds after lid removal, and **(D)** dispersal of the honeycomb pattern 95 seconds after lid removal. Insets are digitally magnified images (2.0x) of the indicated area. The corresponding video can be viewed as Movie S1. This movie is representative of four biological replicates. The diameter of the Petri dish is 10 cm.

**Figure 3:**
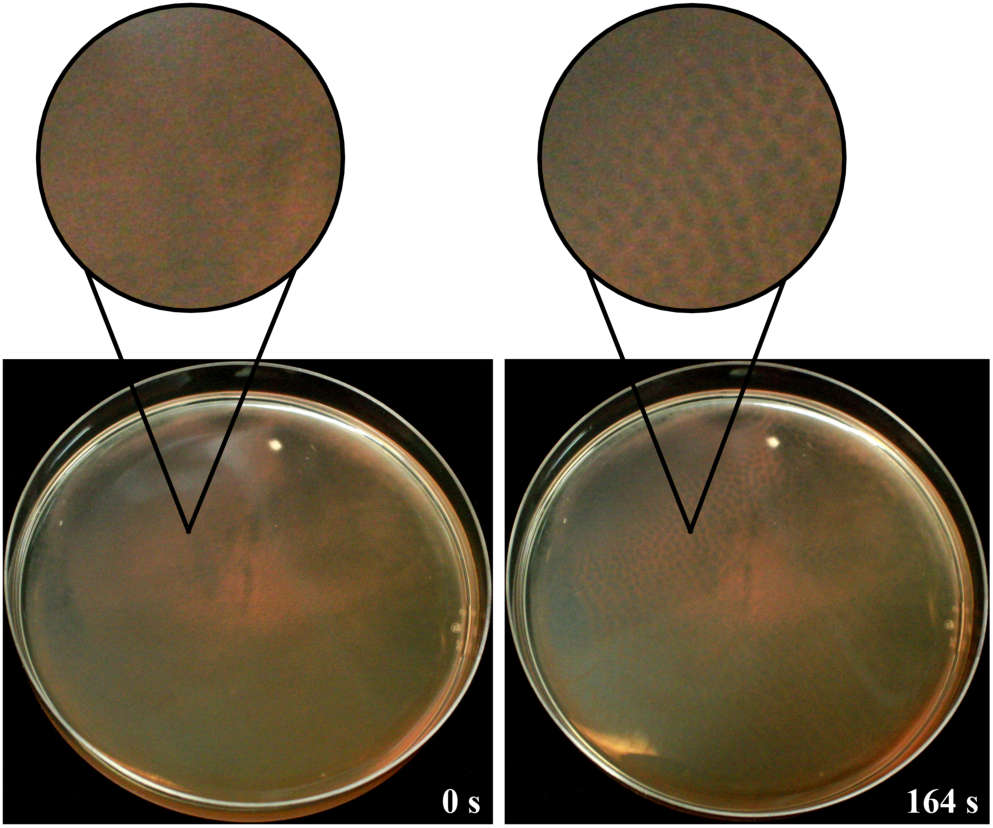
Honeycomb patterns form in the absence of oxygen. In a wildtype culture, an immersed liquid biofilm had formed after 18 hours at room temperature in an anaerobic chamber. After this incubation, the removal of the Petri dish lid resulted in the formation of honeycomb patterns. Representative images were taken immediately after opening the lid (left) and after 2 minutes and 44 seconds (right). Insets are digitally magnified images (2.9x) of the indicated area. Images were taken in suboptimal lighting in the anaerobic chamber; brightness and contrast were adjusted in both images for clarity. Images shown are representative of two replicates tested. The Petri dish diameter is 10 cm.

**Figure 4:**
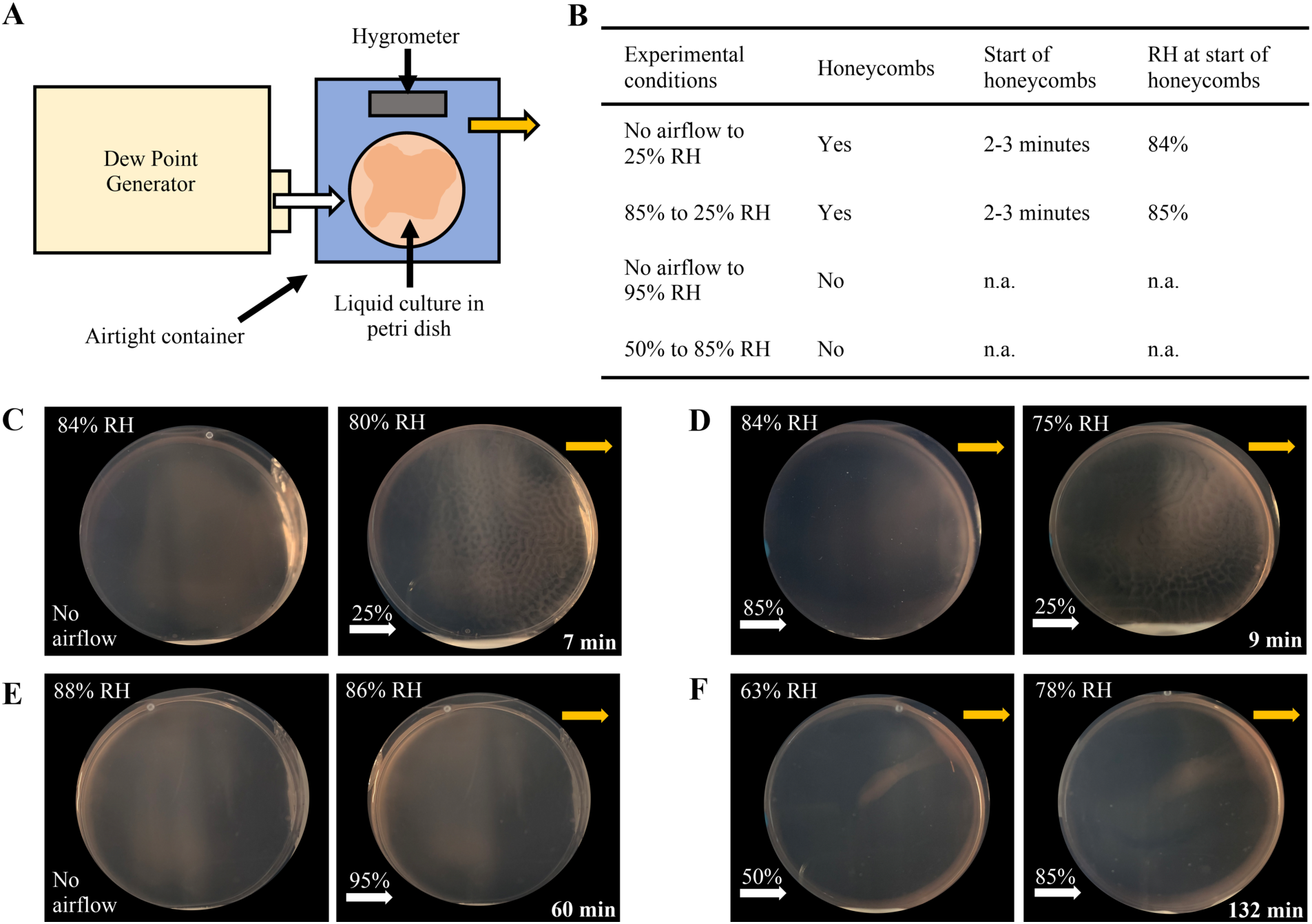
Lowering of humidity levels induces honeycomb pattern formation. **(A)** Schematic representation of the Dew Point Generator (DPG) setup for controlled humidity experiments. The DPG was attached to a small, airtight plastic container with an input airflow tube (white arrow) from the DPG and an output airflow tube (yellow arrow). A lidless Petri dish with liquid culture and a hygrometer were placed inside the container. All experiments were carried out at room temperature. **(B)** Table summarizing results from DPG experiments. ‘Start of honeycombs’ refers to the time at which honeycomb patterns began to form; ‘RH at start of honeycombs’ refers to the RH as measured by the hygrometer inside the container at the same time. **(C, E)** No air was flowing overnight, and after 18 hours, 25% or 95% RH airflow was dispensed from the DPG, respectively. **(D, F)** RH humidity settings were changed from 85% to 25% and 50% to 85% RH, respectively, after an immersed liquid biofilm was formed with constant airflow overnight. In (**C-F)**, images on the left are representative of the immersed liquid biofilm at the start of the experiment and show the RH in the container (top left corner) as measured on the hygrometer at that time. Images on the right are representative of the results obtained over the course of the experiment. For **(C)** and **(D)**, RH and time at the peak of honeycomb pattern formation are shown (top left and bottom right corner, respectively). For **(E)** and **(F)**, RH and time are indicated for when the experiment was concluded. White arrows indicate the entry point of the airflow from the DPG with the above percentage indicating the RH of the input airflow. Yellow arrows indicate the airflow exit point. Experiments in (**C-F)** are representative of at least two biological replicates. The diameter of the Petri dishes is 10 cm.

### Quantification of immersed liquid biofilm coverage

Immersed liquid biofilm coverage within the Petri dish was quantified using Fiji (53) by converting the image to grayscale and binary, drawing a region of interest (ROI) around the Petri dish, and measuring the area corresponding to the biofilm as a percentage of the total ROI. Each strain was tested at least twice.

### Kinetics of honeycomb pattern formation and dispersal

Calculation of when the immersed liquid biofilm began making honeycomb patterns was determined by measuring the time it took for honeycomb patterns to form after the lid of the Petri dish was removed. Time to peak honeycomb formation was defined as the point at which honeycombs were the clearest and covered the greatest extent of the plate after lid removal. The point of dispersal was defined as the time at which honeycombs moved substantially outward, distorting their initial configuration. Each strain was tested at least twice.

### *H. volcanii* anaerobic growth curve

To optimize Hv-Cab medium for anaerobic growth, we tested fumarate concentrations between 0 mM and 60 mM with 25 mM final concentration of PIPES buffer (adapted from (54)). Hv-Cab anaerobic medium used for anaerobic immersed liquid biofilm and honeycomb pattern formation experiments contained 45 mM sodium fumarate (Acros Organics) with 25 mM final concentration of PIPES buffer (Alfa Aesar, 0.5 M, pH 7.5); uracil added to this medium was dissolved in ddH_2_O (Millipore Sigma) (final concentration 2 μg/mL) rather than DMSO. Media was de-gassed in the microaerobic chamber 24 hours before use. *H. volcanii* liquid cultures were inoculated from colonies into 5 mL of each of the six fumarate Hv-Cab media followed by continuous shaking at 45°C. Subsequently, each culture was transferred into wells of a 96-well plate and diluted to an OD_600_ of 0.01 (with the exception of the 0 mM fumarate medium, which was diluted to 0.005) with fresh liquid medium added to bring the final volume to 300 µL (16 technical replicates of one biological replicate). OD_600_ recordings were taken every 30 minutes for 44 hours and then every 60 minutes for 96 hours (6 days total) with an Epoch 2 Microplate Spectrophotometer (Biotek, Winooski, VT) at 45°C within a rigid gloveless hypoxic chamber (Coy Lab, Grass Lake, MI). The plate underwent a double orbital shake for one minute before each measurement.

### Assessment of culture colors in different media

After observing a darker coloration of cultures in anaerobic media, we tested the effects of different media components on the color of *H. volcanii* cultures under aerobic conditions. *H. volcanii* cultures were grown in Hv-Cab medium, Hv-Cab medium with 25 mM PIPES buffer, and Hv-Cab with 25 mM PIPES buffer and 45 mM sodium fumarate. After inoculation and growth at 45°C to an OD_600_ between 0.4 and 0.8, cultures were diluted to an OD_600_ of 0.05 and grown for 48-hours at 45°C. Cultures were then diluted to the same stationary phase OD_600_ and imaged with an iPhone to assess color differences.

### Immersed liquid biofilm and honeycomb pattern formation in anaerobic chamber

Strains were inoculated aerobically in 5 mL of 45 mM fumarate Hv-Cab medium followed by overnight aerobic incubation at 45°C with shaking (orbital shaker at 250 rpm) until the strains reached mid-log phase (OD_600_ 0.3-0.7). Mid-log cultures were diluted to an OD_600_ of 0.05 at a final volume of 20 mL followed by aerobic shaking incubation at 45°C for 48 hours. After the 48 hours incubation period, cultures were poured into sterile Petri dishes (100mm x 15mm, Fisherbrand) in an anaerobic chamber (Coy) with a Palladium catalyst; oxygen gas was purged and replaced with a gas mix of hydrogen/nitrogen (5%/95%). Poured cultures were left in the anaerobic chamber for 24 hours in an incubator (41°C), after which the resulting immersed liquid biofilm and honeycomb pattern formation were observed and imaged. Note that for one of the plates that was tested, the oxygen level in the anaerobic chamber was between 7 and 13 ppm. Strains were left in the anaerobic chamber for an additional 18 hours either at room temperature or in an incubator (45°C) and then observed again for both immersed liquid biofilms and honeycombs.

### Dew Point Generator

Experiments were performed using the same protocol for immersed liquid biofilm formation with the exception that Petri dishes were not covered with Petri dish lids, and the Petri dishes were placed in plastic airtight containers connected to a Dew Point Generator (DPG) (LI-610 Portable Dew Point Generator, LI-COR) at room temperature. Air from the DPG entered the container through a silicone tube and exited the container through a silicone tube at the opposite end of the container. The airflow was dispensed at 16-20 cm^3^/min at the appropriate temperature to confer the desired RH level (calculated as described in the manual). The inside of the airtight container was lined with Styrofoam and aluminum foil to reduce the headspace of the Petri dish and therefore concentrate the distributed airflow. A hygrometer (AcuRite) was also present inside the container to measure RH levels.

## Results

### Development of a rapid immersed liquid biofilm formation assay

Chimileski et al. described the formation and maturation of static liquid biofilms from late-log phase (OD_600_ of 1.0) liquid shaking cultures after an incubation period of seven days (6). To further characterize immersed liquid biofilms and determine the *H. volcanii* proteins required for its formation, we set out to develop a fast and reproducible protocol for immersed liquid biofilm formation. Using a stationary phase liquid culture transferred from a shaking culture tube into a Petri dish, we observed that *H. volcanii* strain H53, the wild-type strain used in this study, begins forming an observable biofilm after as little as eight hours of static incubation, with a robust biofilm being formed within 15 hours and not changing significantly for the next six hours. Therefore, we chose to set our standard immersed liquid biofilm observation time at 18 ± 3 hours of static incubation (Fig. 1A).

While the timing of immersed liquid biofilm formation under the conditions tested was reproducible, they presented stochastic variations in shape, color intensity (likely based on differences in cell density), and coverage of the Petri dish (Fig. 1B). The shape of immersed liquid biofilms in this study ranged from dense, circular areas to diffuse, amorphous shapes. Coverage of the dish varied widely, with the coverage of the area ranging from 67% to 100% in 25 wild-type plates and an average coverage of 91% ± 10% (Fig. S1A).

### Immersed liquid biofilm formation is independent of known H. volcanii components required for biofilm formation on surfaces at the air-liquid interface

Similar to many other archaea and bacteria, evolutionarily-conserved type IV pili are required for *H. volcanii* biofilm formation on surfaces at the air-liquid interface (38, 40). To determine whether type IV pili are also important for immersed liquid biofilm formation, we tested the Δ*pilA1-6* and Δ*pibD* strains, which encode the adhesion pilins and the prepilin peptidase, respectively. Neither of them adhere to coverslips at the air-liquid interface of a liquid culture after 24 hours of incubation (36, 38, 39). Both the *H. volcanii* Δ*pibD* and Δ*pilA1-6* strains formed immersed liquid biofilms comparable to those of wild-type (Table 2 and Fig. S2A). The ability of cells lacking PibD to form these liquid biofilms is particularly notable, as it is responsible for processing all pilins in *H. volcanii* (55). Furthermore, consistent with these results, the Δ*pilB3C3* strain lacking the ATPase (PilB) and the transmembrane component (PilC), both of which are required for PilA1-6 pili assembly (40, 48), as well as the recently characterized Δ*pilB1C1* strain, which lacks PilB and PilC homologs that are likely involved in assembling pili composed of a distinct set of pilins and exhibits defective surface adhesion (41), also formed immersed liquid biofilms similar to those of wild-type. A mutant strain lacking both *pilB* and *pilC* paralogs (Δ*pilB1C1B3C3*) can also form immersed liquid biofilms (Table 2 and Fig. S2A).

**Table 2:** Motility and adhesion mutants lack an immersed liquid biofilm phenotype^*^.

| Strain | Motility<sup>1</sup> | Surface Adhesion<sup>2</sup> | Microcolony Formation<sup>3</sup> | Immersed Liquid Biofilm Formation | References |
| --- | --- | --- | --- | --- | --- |
| H53 | ++ | ++ | ++ | ++ | (38)<sup>1,2</sup>, (40)<sup>3</sup> |
| $\Delta pibD$ | - | - | - | ++ | (38)<sup>1,2,3</sup> |
| $\Delta pilA1-6$ | + | - | - | ++ | (40)<sup>1,2,3</sup> |
| $\Delta pilB1C1$ | n.d. | + | + | ++ | (41)<sup>2,3</sup> |
| $\Delta pilB3C3$ | ++ | + | + | ++ | (40)<sup>1,2</sup>, (41)<sup>3</sup> |
| $\Delta pilB1C1B3C3$ | n.d. | + | + | ++ | (41)<sup>2,3</sup> |
| $cheB::tn$ | - | + | n.d. | ++ | (42)<sup>1,2</sup> |
| $cheF::tn$ | - | + | n.d. | ++ | (Unpublished)<sup>1,2</sup> |
| $\Delta arlA1$ | - | n.d. | n.d. | ++ | (48)<sup>1</sup> |
| $\Delta arlA2$ | +++ | n.d. | n.d. | ++ | (48)<sup>1</sup> |
| $\Delta arlA1-2$ | - | ++ | n.d. | ++ | (38)<sup>1,2</sup> |
| $\Delta aglB$ | - | ++ | ++ / early | ++ | (56)<sup>2,3</sup>, (57)<sup>1</sup> |
| $\Delta agl15$ | n.d. | n.d. | n.d. | ++ | |
| $\Delta pssA$ | + | n.d. | n.d. | ++ | (51)<sup>1</sup> |
| $\Delta pssD$ | + | n.d. | n.d. | ++ | (51)<sup>1</sup> |
| H98 | ++ | ++ | n.d. | ++ | (38)<sup>1,2</sup> |
| $\Delta cetZ1$ | n.d. | n.d. | n.d. | ++ | |

\*Phenotypes are described semi-quantitatively as “-”, no; “+”, reduced; “++”, wild-type like; “+++”, increased motility, surface adhesion or microcolony formation. All tested strains exhibited wild-type like immersed liquid biofilm formation.
<sup>1,2,3</sup> For each column, the reference corresponding to the phenotype is indicated.

Since a screen of an *H. volcanii* transposon insertion library for motility or adhesion-defective *H. volcanii* mutants revealed two mutant strains having insertions in the intergenic regions between chemotaxis genes *cheB* (*hvo_1224*) and *cheW1* (*hvo_1225*) (42) and one mutant strain with a transposon insertion within *cheF* (*hvo_1221*) (data not shown) that have severe motility and adhesion defects, chemotaxis likely plays an important role in adhesion as a prerequisite to biofilm formation in *H. volcanii*. However, immersed liquid biofilm formation comparable to that of *H. volcanii* wild-type cultures was observed in the *cheB::tn* as well as *cheF::tn* mutant strains (Table 2 and Fig. S2A).

As noted, *cheB::tn* and *cheF::tn* are also non-motile, strongly suggesting that archaella – which are required for swimming motility, but, unlike in some other archaea, are not required for biofilm formation on surfaces in *H. volcanii* (38) – are also not involved in immersed liquid biofilm formation in *H. volcanii*. Three archaellin mutants, Δ*arlA1* (nonmotile), Δ*arlA2* (hypermotile), and the double knockout Δ*arlA1-2* (nonmotile), were able to form immersed liquid biofilms comparable to wildtype (Table 2 and Fig. S2A). Strains that lack AglB, the oligosaccharyl-transferase involved in *N*-glycosylation of archaellins and type IV pilins more quickly forms microcolonies compared to wild-type in *H. volcanii* (56). However, neither the Δ*aglB* strain, nor a deletion strain lacking a gene encoding a key component of a second *N*-glycosylation pathway Agl15, conferred immersed liquid biofilm formation defects (Table 2 and Fig. S2A).

We also tested for immersed liquid biofilm formation in deletion mutants involved in lipid anchoring of archaeosortase (ArtA) substrates (51). We speculated that the proper anchoring of some of these ArtA substrates, which includes the S-layer glycoprotein, might potentially be required for immersed liquid biofilm formation. However, two proteins critical for lipid anchoring of ArtA substrates (51), the phosphatidylserine synthase (PssA) and the phosphatidylserine decarboxylase (PssD), do not appear to be required for formation of these liquid biofilms, as the deletion strains, Δ*pssA* and Δ*pssD*, respectively, form immersed liquid biofilms similar to those of the wild-type. Finally, the ability to form rods did not appear to be important for immersed liquid biofilm formation, as the Δ*cetZ1* strain, which lacks the ability to form rods (52), formed these biofilms (Table 2 and Fig. S2A).

Similar to the wild-type strain, immersed liquid biofilms of varying shape and color were formed by the mutant strains tested and by the Δ*cetZ1* parental strain H98 (Fig. S2A). The extent of Petri dish coverage did not differ substantially from the wild-type (Fig. S1A). Overall, these results indicate that key components of the machinery required for surface adhesion, microcolony formation, and swimming motility are not involved in immersed liquid biofilm formation.

### Immersed liquid biofilms self-assemble into honeycomb patterns upon removal of the Petri dish lid

While testing strains for their ability to form immersed liquid biofilms in Petri dishes, we observed a previously undescribed phenomenon: removing the lid of the Petri dish reproducibly caused a rapid, but transient, macroscopic, three-dimensional change in the organization of the immersed liquid biofilm that resulted in the formation of honeycomb-like structures (Fig. 2; Movie S1). After incubation at 45°C for 18 ± 3 hours, removal of the Petri dish lid led to the emergence of a readily observable honeycomb pattern in the immersed liquid biofilm that started within 20 ± 4 seconds (range: 13 s to 27 s) after lid removal in the wild-type strain (Fig. S1B). When the immersed liquid biofilm was incubated at room temperature, the honeycomb pattern emerged more slowly, as honeycombs began to form 92 ± 8 seconds after lid removal on average (range: 81 s to 103 s) (data not shown).

The pattern generally begins in one to two sections of the dish and quickly spreads to cover the biofilm until it reaches its peak formation (Fig. 2B); in wild-type, peak honeycomb formation occurred 38 ± 7 seconds (range: 25 s to 55 s) after lid removal (Fig. S1C). The honeycomb patterns are transient, as dissipation of the honeycombs begins 29 ± 9 seconds (range: 18 s to 57 s) after the peak of honeycomb pattern formation in wild-type (Fig. 2C; Fig. S1D). Interestingly, while the immersed liquid biofilms form close to the bottom of the Petri dish, the hon-eycomb-like structures extend further into the liquid and appear to dissipate close to the ALI (Movie S2). After honeycomb pattern formation and subsequent dissipation, placing the lid back onto the plate and allowing the immersed liquid biofilm to reform for at least one hour enables the pattern formation to again occur once the Petri dish lid is removed again. Honeycomb pattern formation is not dependent on light, as removing the lid in a dark room results in honeycombs as well (data not shown).

The formation of honeycomb patterns can be split into four distinct phases: pre-honeycomb pattern formation, which consists of the immersed liquid biofilm before honeycomb pattern formation begins (Fig. 2A); start of honeycomb pattern formation, when the first honeycombs appear (Fig. 2B); peak-honeycomb pattern formation, which is when the honeycomb pattern is clearest and covers the greatest extent of the biofilm (Fig. 2C); and dispersal of honeycomb patterns, which occurs when the honeycomb pattern begins to dissipate and eventually returns to the settled biofilm state (Fig. 2D). Similar to our results showing that each mutant strain tested was able to form an immersed liquid biofilm, every mutant strain tested also formed honeycomb patterns (Fig. S2B) and honeycomb pattern formation followed a similar time frame compared to that of wild-type in all three phases (Fig. S1B, C, and D).

### Honeycomb pattern formations occur under anaerobic conditions

To determine the factor(s) that induce the morphological change upon removal of the lid, we next sought to identify conditions under which honeycomb-like structures fail to develop. Having determined that honeycomb patterns were observed in Petri dishes as well as 6- and 24-well plates but not in standing tubes containing 5 mL liquid cultures (data not shown), we first investigated whether differences in oxygen concentrations play a role in honeycomb pattern formation. While *H. volcanii* is a facultative anaerobe, to the best of our knowledge, *H. volcanii* biofilm experiments had not previously been carried out under anaerobic conditions.

Following a previous study (54), we modified the Hv-Cab medium to contain fumarate as an electron acceptor and PIPES as a buffer. We tested a range of fumarate concentrations along with 25 mM of PIPES buffer via an anaerobic growth curve using a 96-well plate assay (Fig. S3A). While fumarate was required for cell growth under anaerobic conditions, differences between the tested fumarate concentrations were neglectable. Therefore, we chose the intermediate concentration of 45 mM fumarate for further experiments (Fig. S3B). Interestingly, we noticed that wild-type cultures grown with 25 mM PIPES and 45 mM fumarate were darker pink in color than those without. To distinguish the effects of different media components, we grew wild-type cultures with Hv-Cab, Hv-Cab with 25 mM PIPES, and Hv-Cab with 25 mM of PIPES and 45 mM of fumarate under aerobic conditions and compared the color at the same stationary phase OD_600_ values. We found that cultures with just PIPES and with both PIPES and fumarate both produced darker cultures relative to the cultures without PIPES, indicating that the presence of the PIPES buffer stimulated higher expression levels of bacterioruberin, the most prevalent carotenoid in *H. volcanii* (Fig. S3C) (58). We also observed that cultures grown with PIPES and fumarate grew to a higher final OD_600_ than cultures without fumarate (data not shown).

Using the fumarate Hv-Cab medium, we tested the ability of wild-type cells to form honeycomb patterns under anaerobic conditions. We used the same protocol as shown in Fig. 1A, with the exception that stationary phase liquid cultures were poured and incubated (at 41°C) in Petri dishes that were maintained in an anaerobic chamber. After 24 hours, immersed liquid biofilms were tested for their ability to form honeycomb patterns by opening the Petri dish lid inside the anaerobic chamber. Interestingly, we determined that the formation of honeycomb patterns under anaerobic conditions was comparable to those observed in aerobic cultures (Fig. 3). The cultures were incubated for an additional 18 hours at either room temperature or at 45°C in the anaerobic chamber, after which the immersed liquid biofilms had re-formed; honeycomb patterns formed upon removal of the lid at the same rates as they did in aerobic cultures at both of these temperatures (Fig. 3).

### Decreasing humidity triggers honeycomb pattern formation

Given that neither light nor oxygen exposure changes caused honeycomb pattern formation, we considered two possibilities: (i) the dissipation of accumulated volatiles triggers honeycomb pattern formation, or (ii) changes in humidity levels lead to the formation of honeycomb-like structures. To distinguish between these two hypotheses, we used a Dew Point Generator (DPG), which dispenses air at controlled humidity levels (Fig. 4A). We attached airtight containers, each containing a liquid culture in a Petri dish lacking a lid as well as a hygrometer to measure RH levels, to the DPG.

We first used the DPG to test whether going from no airflow to 25% RH airflow would cause honeycombs to form. This experiment is akin to the original lid-lifting experiment. The experimental setup with no airflow allowed humidity levels inside the container to reach percentages ranging from the low to upper 80s. It also enabled any potential volatiles to accumulate. We observed that a diffuse immersed liquid biofilm had formed after 18 hours, and switching on the airflow at 25% RH triggered honeycomb pattern formation (Fig. 4C). While this experiment cannot distinguish between the humidity change and volatile dissipation hypotheses, it showed that we can replicate the effect of lifting the lid within a setting with controlled airflow.

Next, instead of letting a humidity level of 85% RH be reached via evaporation of the culture, we generated this level of humidity by dispensing 85% RH airflow for 18 hours post-pouring, upon which a diffuse immersed liquid biofilm had formed. In this setup, the humidity level is unaffected, but the potential accumulation of volatiles is prevented. The humidity of the airflow was then changed to 25% RH while maintaining the same flow rate. The DPG requires several minutes to reach the new set humidity level; when the DPG was dispensing air at about 60% RH, honeycomb patterns were triggered to form (approximately 2-3 minutes following the initial switch from 85% RH to 25% RH) and took about 10 minutes to cover the entire plate (Fig. 4D). This result indicates that an accumulation of volatiles in the headspace, prevented by the constant airflow during the 18-hour incubation time, was neither required for immersed liquid biofilm nor for honeycomb pattern formation. In addition, in a separate experiment, 95% RH airflow did not trigger honeycomb pattern formation in immersed liquid biofilms that were allowed to form overnight with no airflow (Fig. 4E). The ability to form honeycomb-like structures was confirmed for these cultures by opening the container lid (data not shown). These results suggest that honeycomb pattern formation is not triggered by the dissipation of volatiles nor by high humidity airflow; instead, a decrease in the humidity level results in honeycomb pattern formation.

This hypothesis is further strengthened by an experimental setup in which the airflow was set to 50% RH overnight, which resulted in a hygrometer reading of 63% RH in the chamber after 18 hours. The humidity level was then increased to 85% RH. Honeycombs did not form even after over an hour of observation (Fig. 4F). However, it should be noted that in this experiment the immersed liquid biofilm had mostly formed along the edges of the Petri dish, which might have precluded our ability to observe honeycombs, should they have been triggered to form. In general, we noticed that liquid biofilms predominantly formed along the edges of the petri dish when lower-level humidity airflow was passed over the plate (Fig. S4A). Conversely, when higher-humidity air was dispensed over the Petri dish, the immersed liquid biofilm that formed was diffuse (Fig. S4B).

## Discussion

In this study, we developed an optimized workflow to observe the development of *H. volcanii* immersed liquid biofilms. Using that workflow, we determined that, for immersed liquid biofilm formation, this model haloarchaeon does not require any of the genes known to affect biofilm formation on abiotic surfaces. Deletion mutants lacking *pilA1-6*, which encode type IV pilins, or *pilB1C1* and *pilB3C3*, which encode proteins required for pilus assembly, all of which exhibit adhesion defects in the ALI assay, could still form immersed liquid biofilms. While the *H. volcanii* genome encodes two additional PilB and PilC paralogs and 36 additional predicted pilins (39), which can presumably form distinct type IV pili, it is unlikely that these proteins are involved in immersed liquid biofilm formation since the absence of *pibD*, which encodes the only *H. volcanii* prepilin peptidase (55) and is required to process prepilins prior to pilus assembly (57), did not affect immersed liquid biofilm formation.

Furthermore, transposon mutants affecting *H. volcanii* chemotaxis genes, which result in decreased ALI adhesion, still exhibited immersed liquid biofilm formation. However, little is known about chemotaxis and intracellular signaling of *H. volcanii*. Thus, it is possible that an alternative signaling pathway is required for the formation of immersed liquid biofilms. Similarly, immersed liquid biofilms formed independent of two major post-translational modifications of cell surface proteins, *N*-glycosylation (Δ*aglB* and Δ*agl15*) and ArtA-dependent C-terminal lipid anchoring (Δ*pssA* and Δ*pssD*). These modifications affect the function of various secreted proteins, including the S-layer glycoprotein. However, that does not preclude that other cell surface proteins are involved in the formation of immersed liquid biofilms.

It is intriguing that none of the genes known to affect adhesion to abiotic surfaces led to an altered immersed liquid biofilm phenotype. The process through which this type of biofilm forms remains to be elucidated. However, during this study, we also observed a previously undescribed phenomenon that could provide further insights into immersed liquid biofilms: the rapid, transient, and reproducible honeycomb pattern formation that occurs in cultures with established immersed liquid biofilms upon removal of the Petri dish lid. Chimileski et al. previously noted the dynamic nature of immersed liquid biofilms; however, their work focused on filamentous structures extending and retracting on the edge of the Petri dish over the course of hours (6). While the time frame of these movements is quite different from the rapid formation of honeycomb patterns described here, they were triggered similarly by what was described as physical agitation (tapping or slight lifting of the Petri dish lid). In fact, while not discussed in the paper, faint honeycomb-like patterns are visible by slowing down a supplemental video by Chimileski et al. (capturing 90 minutes in 10 second intervals) (6). As we describe here, after incubation at 45°C, honeycomb patterns formed on average within 20 ± 4 seconds after lid removal and dissipated on average after 67 ± 13 seconds after lid removal (corresponding to 29 ± 9 seconds after peak honeycomb pattern formation). Therefore, it is likely that the social motility discussed by Chimileski et al. represents the subsequent events following the rapid honeycomb pattern formation described here.

Since we showed here that honeycomb-like structures formed rapidly even in non-motile and non-piliated mutants, together with the short time frame of honeycomb formation, our results strongly suggest that this process is not driven by active movement of cells. The short time frame of honeycomb pattern formation also indicates that whatever is passively moving the cells must be present within the immersed liquid biofilm before honeycombs form. Therefore, honeycomb pattern formation may reveal the underlying structure of the immersed liquid biofilm. It has been hypothesized that the EPS components of an *H. volcanii* immersed liquid biofilm include, primarily, polysaccharides, eDNA, and amyloid proteins (6). These EPS components likely form the underlying structure providing support for the biofilm, and under the conditions tested in this study, this skeletal structure may have played a direct role in the formation of honeycomb patterns. While EPS biosynthesis pathways in *H. volcanii* remain to be characterized, the pathway of exopolysaccharide biosynthesis in *H. mediterranei* has been determined (59). Interestingly, both immersed liquid biofilm formation and honeycomb pattern formation also occurred in *H. mediterranei* (Fig. S5), suggesting that the genes required for both processes are conserved between these species.

While, to the best of our knowledge, the rapid transition from diffuse immersed liquid biofilms into honeycomb patterns has not been described so far, honeycomb-like structures have been observed in biofilms of other prokaryotes. These honeycomb patterns often appear to serve structural roles within the biofilm and form on a microscopic scale (diameters of five to 50 µm compared to one to five mm of *H. volcanii* honeycomb like structures described in this study) over the course of hours to days (i.e. multiple generation times). For example, a honeycomb-like meshwork generated by interconnected eDNA strands bound to cells through positively charged proteins has been reported for *Staphylococcus aureus* biofilms (60), and membrane-bound lipoproteins that can bind DNA have been implicated in maintaining the structure of *S. aureus* biofilms (61). Furthermore, in *P. aeruginosa* PAO1 biofilms, interactions between eDNA and exopolysaccharide Psl fibers result in web-like patterns observed in pellicles and flow-cells (62, 63). The web pattern might function as a supportive scaffold that allows bacterial attachment and subsequent growth within the biofilm (63). Furthermore, it might play a role in bacterial migration to facilitate nutrient uptake, since the web-like pattern is most pronounced in nutrient-starved areas within the biofilm (62). This is in line with studies in *Listeria monocytogenes* biofilms which, under conditions of constant liquid flow, form honeycomb-like (‘knitted’) structures in diluted, nutrient-poor media but not in rich media (64); under static conditions, honeycomb hollows were shown to contain planktonic cells, perhaps suggesting a transition to biofilm dispersal (65). A variety of benefits from honeycomb-like structures is also supported by Schaudinn *et al*., who hypothesize that for cells undergoing stress from fluid forces, honeycombs could provide flexibility and distribution of forces over the six vertices (66). Moreover, the increased surface area of honeycombs-like structures could aid cells faced with limited nutrients and could potentially also serve as communication ‘roadways’ for inter-cellular signaling (66). Computer models of honeycomb patterns with a larger diameter (several hundred µm) observed in *Thiovulum majus* biofilms suggest that these structures cause water advection that would result in improved distribution of oxygen within the biofilm (67).

While the microscopic dimensions of these bacterial honeycomb patterns are substantially different from the macroscopic scale of the honeycomb patterns we described here for *H. volcanii*, these examples illustrate that honeycomb-like structures may serve important biological roles. Upon honeycomb pattern formation, an upward motion (towards the ALI) of cells was observed, followed by the dissipation of the pattern (Movie S2). Since an active movement of cells is unlikely due to the lack of flagella and type IV pili in the respective mutants, which still formed honeycomb patterns, the honeycomb-like structures might contribute to increased floating properties of the biofilm. Based on the rapid formation of honeycomb patterns in *H. volcanii*, which is unparalleled in other prokaryotes, it is also tempting to speculate that this process results in turbulences in the liquid culture that could facilitate improved distribution of minerals and other nutrients from the surrounding media to the cells within the biofilm.

The DPG experiments with constant airflow indicate that the rapid transition to honeycomb-like structures was not induced by changes in the concentration of volatiles synthesized by *H. volcanii*. Instead, decreasing the humidity level within the headspace of the immersed liquid biofilm triggered honeycomb pattern formation. Biofilm formation has previously been shown to be influenced by RH levels (35). Moreover, in *B. subtilis*, expansion of biofilm coverage area was observed with increases from low (20-30%) to high (80-90%) RH levels (68). Others have shown via simulation models that phase separation can occur within a biofilm due to aggregation of bacterial cells to provide ample volume for produced EPS (31) or as a result of cell-cell and cell-surface interactions (32). While these models focused on microscopic pattern formations, it is nevertheless tempting to speculate that in *H. volcanii* the reduction in humidity led to phase separation resulting in honeycomb pattern formation. However, this does not exclude the possibility that the observed honeycomb-like structures provide a fitness benefit to the organism. Honeycombs may protect against increased evaporation at lower humidity levels: in general, pattern formation has been suggested to confer cells within biofilms increased protection against environmental flux (69), potentially extending to changes in humidity. EPS has also been suggested to be a protective measure against changing external conditions such as humidity (70). Alternatively, in the natural environment of *H. volcanii*, humidity dispersed by wind could signal a beneficial change in environmental conditions, e.g. the mixing of water or an influx of oxygen, and honeycomb pattern formation may aid in the dispersal of immersed liquid biofilms. This hypothesis is supported by the outward and upward movement of cells following honeycomb pattern formation. Similar to the formation of immersed liquid biofilms, the molecular mechanism and genes required to form honeycomb-like structures in *H. volcanii* remain to be elucidated. This could provide further insights into the biological role of these structures as well.

In conclusion, this study showed that *H. volcanii* immersed liquid biofilms form through an as-of-yet-unknown mechanism that is independent of many of the genes required for biofilm formation at the ALI. Moreover, this study supports the notion that pattern formation within biofilms is a common phenomenon, but in contrast to previously described pattern formations in bacteria, honeycomb-like structures in *H. volcanii* can form on a macroscopic scale and within seconds, triggered by reduction in humidity levels.

## Acknowledgments

We would like to thank Fevzi Daldal, Elliot Friedman, Mark Goulian, John Hallsworth, Brent Helliker, Chinedum Osuji, Friedhelm Pfeiffer, and Gary Wu for helpful discussions and help with experiments. We acknowledge the Microbial Culture & Metabolomics Core of the PennCHOP Microbiome Program for assistance with culture studies. We also thank the Daldal and Helliker labs for the use of the anaerobic chamber and Dew Point Generator, respectively, as well as Ian Duggin for providing the Δ*cetZ1* strain.

S.S. was supported by the German Research Foundation (DFG Postdoctoral Fellowship, 398625447). M.P., H.S. and Z.M. were supported by the National Science Foundation Grant 1817518. C.d.V., C.R. and J.S. were supported by the Dept. of Biology Biol376 lab course fund. A.W.B.F. was supported by his personal startup fund from Brandeis University. The funders had no role in study design, data collection and interpretation, or the decision to submit the work for publication.

## Supplemental Material

**Figure S1: Immersed liquid biofilms of all analyzed mutant strains cover a similar Petri dish area and exhibit similar timing in their formation and honeycomb patterns as the wild-type**. Boxplots for all analyzed strains represent the area of a Petri dish covered by the immersed liquid biofilm (ILB) **(A)**, the time to the start of honeycomb pattern (HCP) formation after lid removal **(B)**, the time to the peak of honeycomb pattern formation after lid removal **(C)**, and the time to dispersal after peak honeycomb pattern formation **(D)**. Box center line, median; box limits, upper and lower quartiles; whiskers, 1.5x interquartile range; points, all individual values. Representative images for all strains can be found in Fig. S2.

**Figure S2: All tested mutant strains formed immersed liquid biofilms as well as honeycomb pattern formations**. Each mutant strain (as well as the parental strains H53 and H98) was tested for **(A)** immersed liquid biofilm formation using the optimized protocol (see Fig. 1) and for **(B)** honeycomb pattern formation after opening the Petri dish lid under aerobic conditions. Images for **(A)** were taken within 10 seconds of Petri dish lid removal, and images for **(B)** were taken at peak honeycomb formation (time to reach peak honeycomb formation from lid removal noted in image). Images shown are representative for at least two replicates tested for immersed liquid biofilm and honeycomb pattern formations for each strain. All strains were tested after incubation at 45°C. The diameter of the Petri dishes is 10 cm. Quantitative analyses for all strains can be found in Fig. S1.

**Figure S3: The addition of fumarate to Hv-Cab allows for growth under anaerobic conditions. (A)** Different fumarate concentrations in Hv-Cab containing PIPES buffer were tested for growth under anaerobic conditions by measuring OD_600_ over six days in a 96-well plate at 45°C. The anaerobic growth curves represent the mean ± SD of 16 technical replicates. **(B)** The difference in OD_600_ between the last and first time point is given as the mean ± SD for the different fumarate concentrations. Only the growth of wildtype cells in medium containing no fumarate was statistically significantly different from growth in all other fumarate concentrations (p < 1e-15). **(C)** Wild-type cultures grown in the presence of PIPES and PIPES with fumarate (middle, right) were darker in color than those grown without PIPES (left). Cultures were diluted to all be at the same OD_600_; those shown here are representative of two biological replicates.

**Figure S4: Immersed liquid biofilm formation differs between low and high RH levels. (A)** After 18 hours of 50% RH airflow, a more condensed immersed liquid biofilm formed mostly at the edges of the Petri dish. **(B)** After 18 hours of 85% RH airflow, a diffuse immersed liquid biofilm can be seen in the center and at the edges of the Petri dish. Arrows represent input airflow tubes (white) and output airflow tubes (yellow). For both **(A)** and **(B)**, at least two replicates were tested. The diameter of the Petri dishes is 10 cm. Experiments were carried out at room temperature.

**Figure S5: *H. mediterranei* forms liquid biofilms and honeycomb patterns. (A)** Representative images of a wild-type immersed liquid biofilm immediately after Petri dish lid removal, followed by **(B)** start of honeycomb formation 14 seconds after lid removal, **(C)** peak honeycomb pattern formation 26 seconds after lid removal, and **(D)** dispersal of the honeycomb pattern 71 seconds after lid removal, are shown. Cultures were incubated at 45°C prior to testing. Insets are digitally magnified images (2.0x) of the indicated area. The Petri dish diameter is 10 cm.

**Movie S1: *H. volcanii* form honeycomb patterns**. Removal of the Petri dish lid, after an immersed liquid biofilm has formed, triggers honeycombs to form and then dissipate. Movie begins 2 seconds after lid removal. Time lapse was acquired at 150 frames per second and played at actual real-time speed. Honeycomb pattern formation begins at 25 seconds and dissipation begins around 40 seconds. Cultures were incubated at 45°C prior to testing. The Petri dish diameter is 10 cm.

**Movie S2: Honeycomb pattern formations extend out- and upwards**. Honeycomb-like structures extend out- and upwards into the liquid and appear to dissipate close to the ALI. Time lapse was acquired at 150 frames per second and played at 15x the actual speed. (Right Panel) 2.5x zoom-in projection of the delimited square on the right panel. Cultures were incubated at 45°C prior to testing. The Petri dish diameter is 10 cm..

