## Supplemental Material for "*Haloferax volcanii* immersed liquid biofilms develop independently of known biofilm machineries and exhibit rapid honeycomb pattern formation"

**This document includes:**

Figures S1 to S5
Captions for Movie S1 and S2

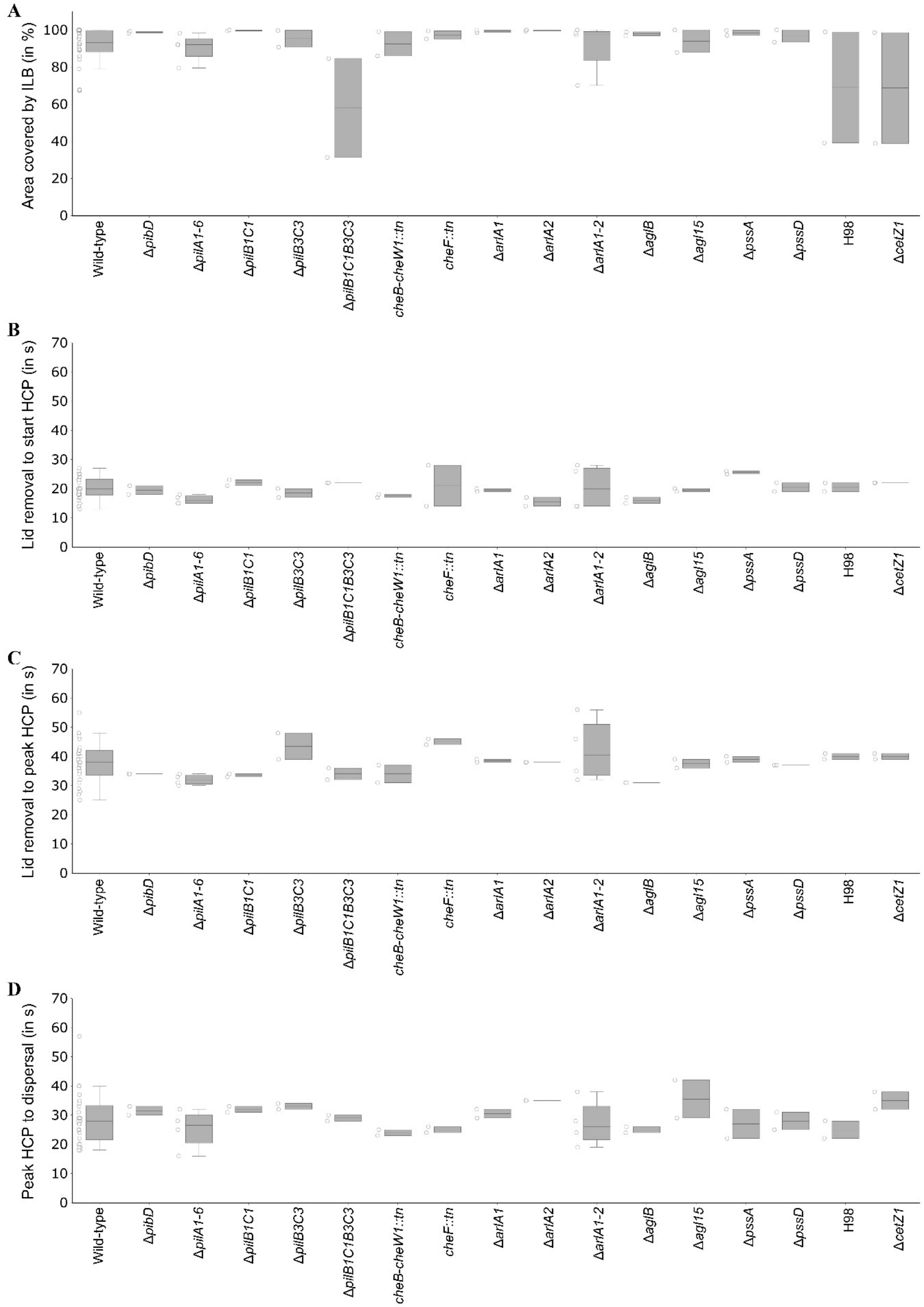


**Figure S1: Immersed liquid biofilms of all analyzed mutant strains cover a similar Petri dish area and exhibit similar timing in their formation and honeycomb patterns as the wild-type.**

Boxplots for all analyzed strains represent the area of a Petri dish covered by the immersed liquid biofilm (ILB) **(A)**, the time to the start of honeycomb pattern (HCP) formation after lid removal **(B)**, the time to the peak of honeycomb pattern formation after lid removal **(C)**, and the time to dispersal after peak honeycomb pattern formation **(D)**. Box center line, median; box limits, upper and lower quartiles; whiskers, 1.5x interquartile range; points, all individual values. Representative images for all strains can be found in Fig. S2.

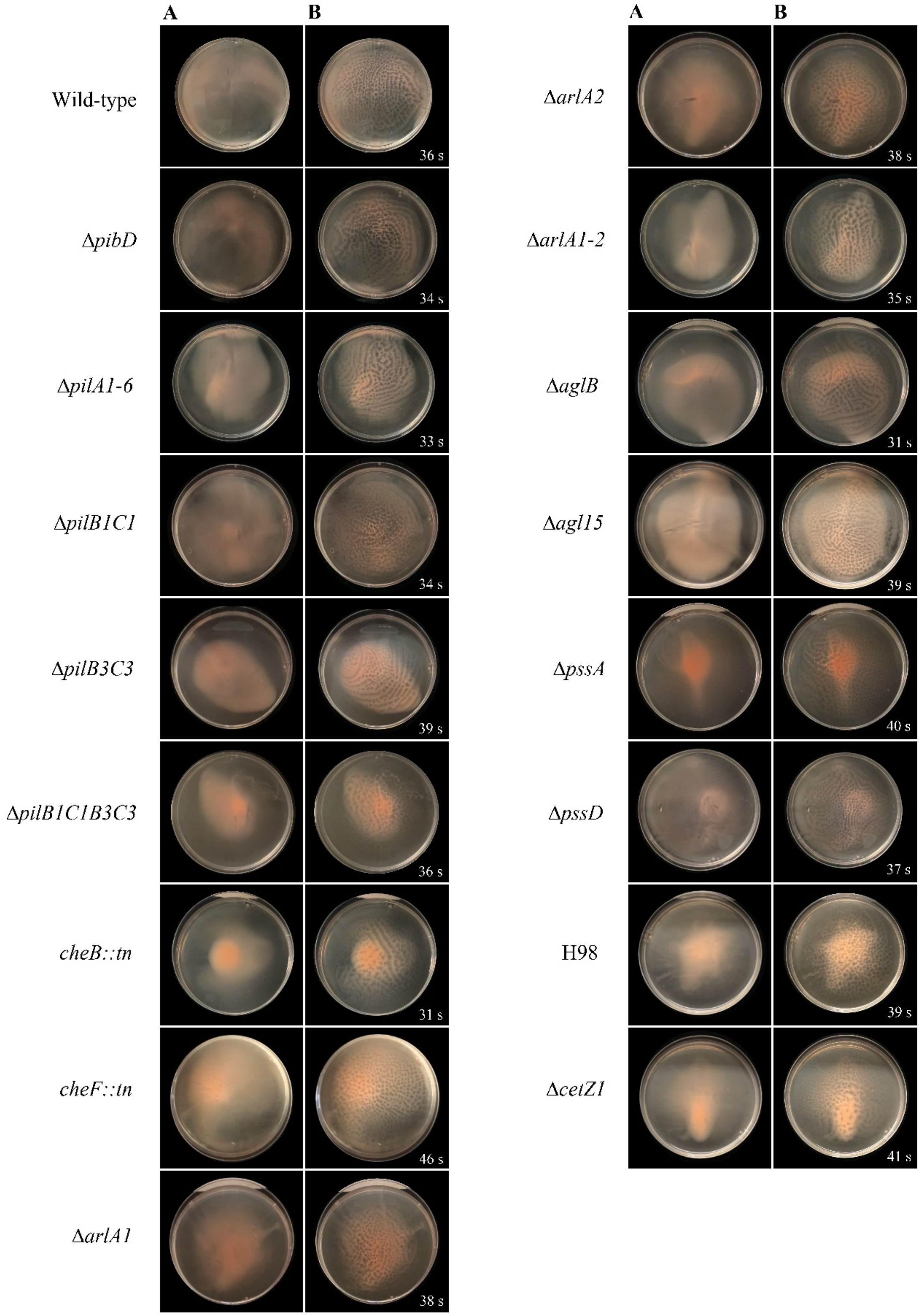


**Figure S2: All tested mutant strains formed immersed liquid biofilms as well as honeycomb pattern formations.**

Each mutant strain (as well as the parental strains H53 and H98) was tested for **(A)** immersed liquid biofilm formation using the optimized protocol (see Fig. 1) and for **(B)** honeycomb pattern formation after opening the Petri dish lid under aerobic conditions. Images for **(A)** were taken within 10 seconds of Petri dish lid removal, and images for **(B)** were taken at peak honeycomb formation (time to reach peak honeycomb formation from lid removal noted in image). Images shown are representative for at least two replicates tested for immersed liquid biofilm and honeycomb pattern formations for each strain. All strains were tested after incubation at 45℃. The diameter of the Petri dishes is 10 cm. Quantitative analyses for all strains can be found in Fig. S1.

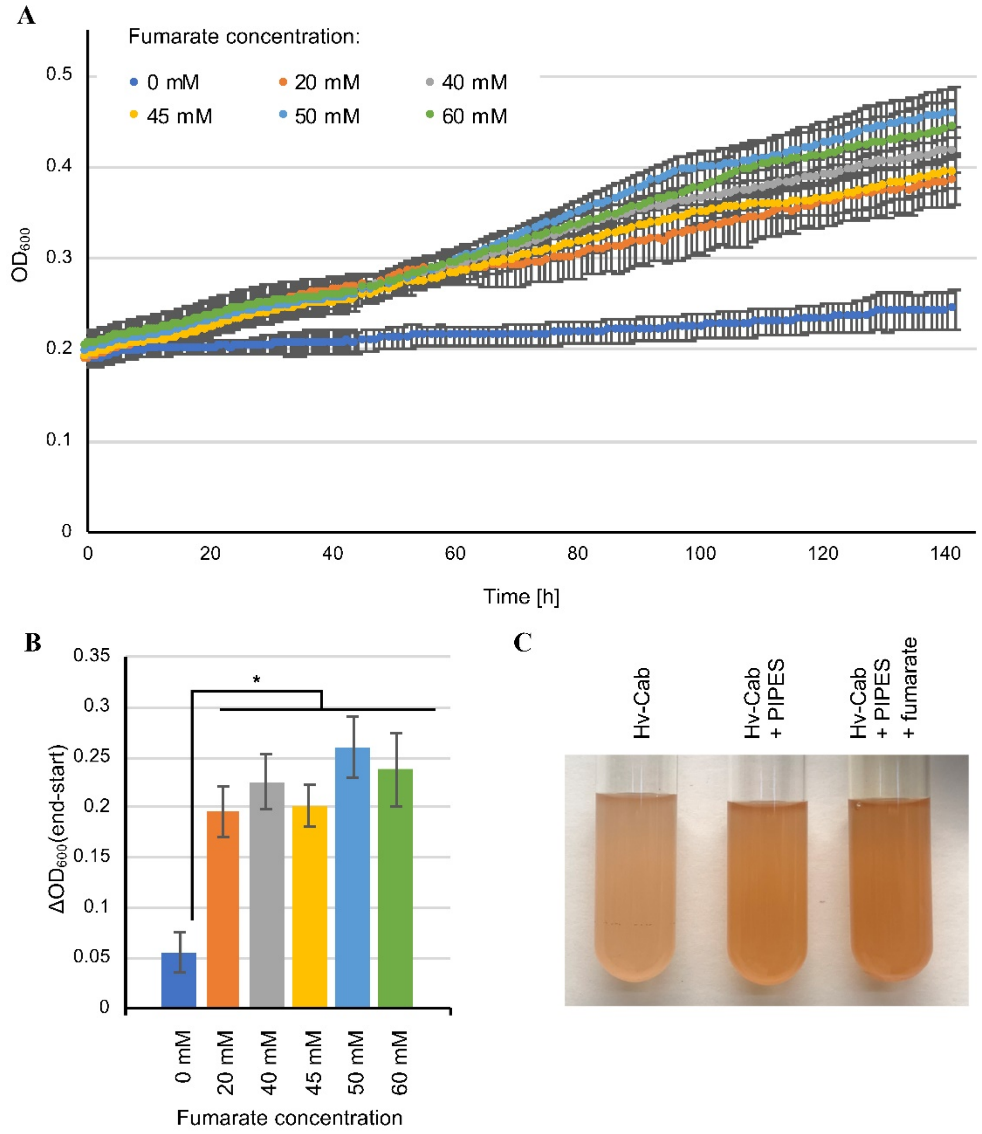


**Figure S3: The addition of fumarate to Hv-Cab allows for growth under anaerobic conditions.** **(A)** Different fumarate concentrations in Hv-Cab containing PIPES buffer were tested for growth under anaerobic conditions by measuring $OD_{600}$ over six days in a 96-well plate at 45°C. The anaerobic growth curves represent the mean ± SD of 16 technical replicates. **(B)** The difference in $OD_{600}$ between the last and first time point is given as the mean ± SD for the different fumarate concentrations. Only the growth of wild-type cells in medium containing no fumarate was statistically significantly different from growth in all other fumarate concentrations ($p < 1e\text{-}15$). **(C)** Wild-type cultures grown in the presence of 25mM PIPES (middle) and 25 mM PIPES with 45 mM fumarate (right) were darker in color than those grown without PIPES (left). Cultures were diluted to all be at the same $OD_{600}$; those shown here are representative of two biological replicates.

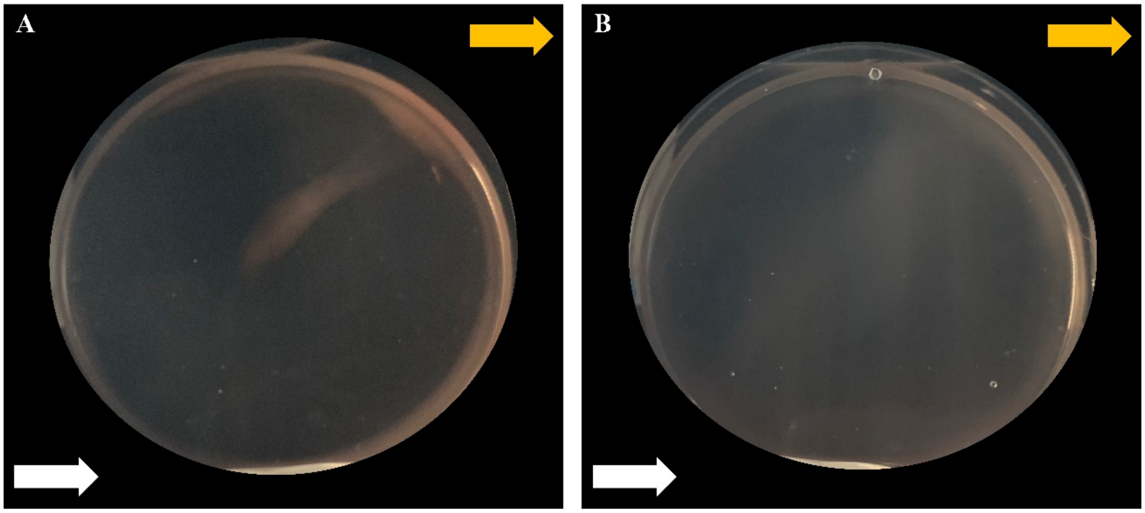


**Figure S4: Immersed liquid biofilm formation differs between low and high RH levels.**
**(A)** After 18 hours of 50% RH airflow, a more condensed immersed liquid biofilm formed mostly at the edges of the Petri dish. **(B)** After 18 hours of 85% RH airflow, a diffuse immersed liquid biofilm can be seen in the center and at the edges of the Petri dish. Arrows represent input airflow tubes (white) and output airflow tubes (yellow). For both **(A)** and **(B)**, at least two replicates were tested. The diameter of the Petri dishes is 10 cm. Experiments were carried out at room temperature.

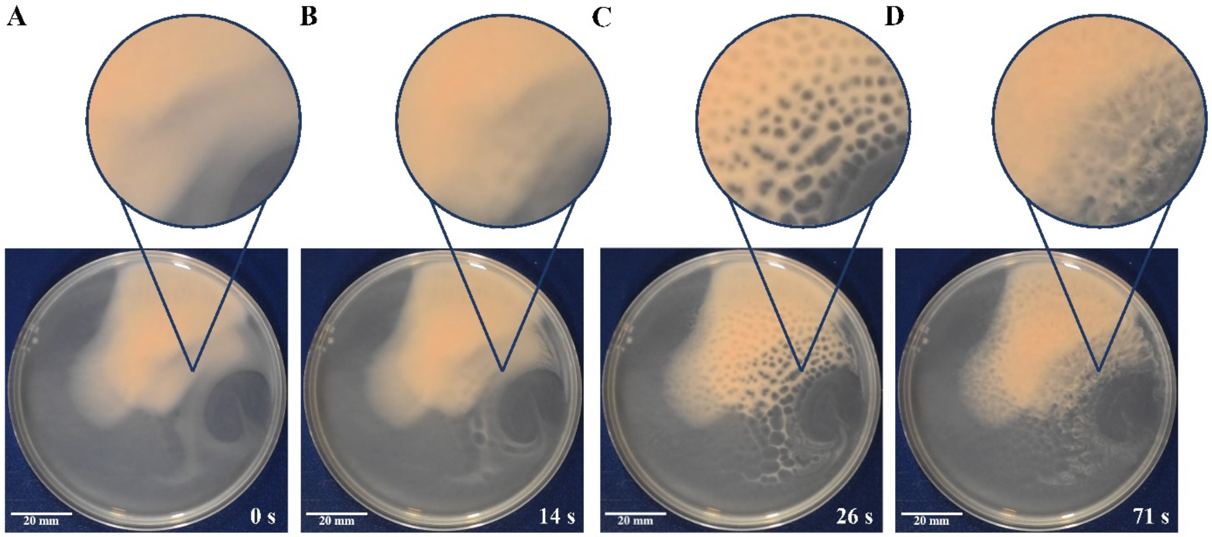


**Figure S5:** ***H. mediterranei* forms liquid biofilms and honeycomb patterns.**
**(A)** Representative images of a wild-type immersed liquid biofilm immediately after Petri dish lid removal, followed by **(B)** start of honeycomb formation 14 seconds after lid removal, **(C)** peak honeycomb pattern formation 26 seconds after lid removal, and **(D)** dispersal of the honeycomb pattern 71 seconds after lid removal, are shown. Cultures were incubated at 45°C prior to testing. Insets are digitally magnified images (2.0x) of the indicated area. The Petri dish diameter is 10 cm.

**Movie S1:** ***H. volcanii*** **form honeycomb patterns.**
Removal of the Petri dish lid, after an immersed liquid biofilm has formed, triggers honeycombs to form and then dissipate. Movie begins 2 seconds after lid removal. Time lapse was acquired at 150 frames per second and played at actual real-time speed. Honeycomb pattern formation begins at 25 seconds and dissipation begins around 40 seconds. Cultures were incubated at 45℃ prior to testing. The Petri dish diameter is 10 cm.

**Movie S2: Honeycomb pattern formations extend out- and upwards.**
Honeycomb-like structures extend out- and upwards into the liquid and appear to dissipate close to the ALI. Time lapse was acquired at 150 frames per second and played at 15x the actual speed. (Right Panel) 2.5x zoom-in projection of the delimited square on the right panel. Cultures were incubated at 45℃ prior to testing. The Petri dish diameter is 10 cm.
